# Interplay between BET Proteins and the Pre-Replication Complex Unveils Therapeutic Vulnerabilities in Cancer

**DOI:** 10.1101/2025.03.27.645172

**Authors:** Kai Doberstein, Vivica Messner, Sebastian Berlit, Benjamin Tuschy, Marc Sutterlin, Frederik Marme

## Abstract

Replication stress is a critical event in cancer development and understanding the underlying molecular mechanisms can help to identify new treatment strategies. Here, we investigate the molecular interplay between the pre-replication complex (pre-RC) and the bromodomain and extraterminal domain (BET) proteins in orchestrating replication stress response. Our findings reveal a mutual dependency between BET proteins and the pre-RC within cancer cells. Notably, reduction in origin licensing or replication initiation makes cells more susceptible to the BET-inhibitor AZD5153. Furthermore, we observe synergistic effects when combining the CDC7 inhibitor XL413 with AZD5153, effects which were partly dependent on *TP53* mutation status. This combination treatment results in unresolved DNA damage and R-loop formation, leading to increased genomic instability. These insights pave the way for novel strategies for targeted cancer therapies and provide a foundation for further investigation into the interplay between CDC7 and BET proteins in replication stress response.

**Graphical abstract:** 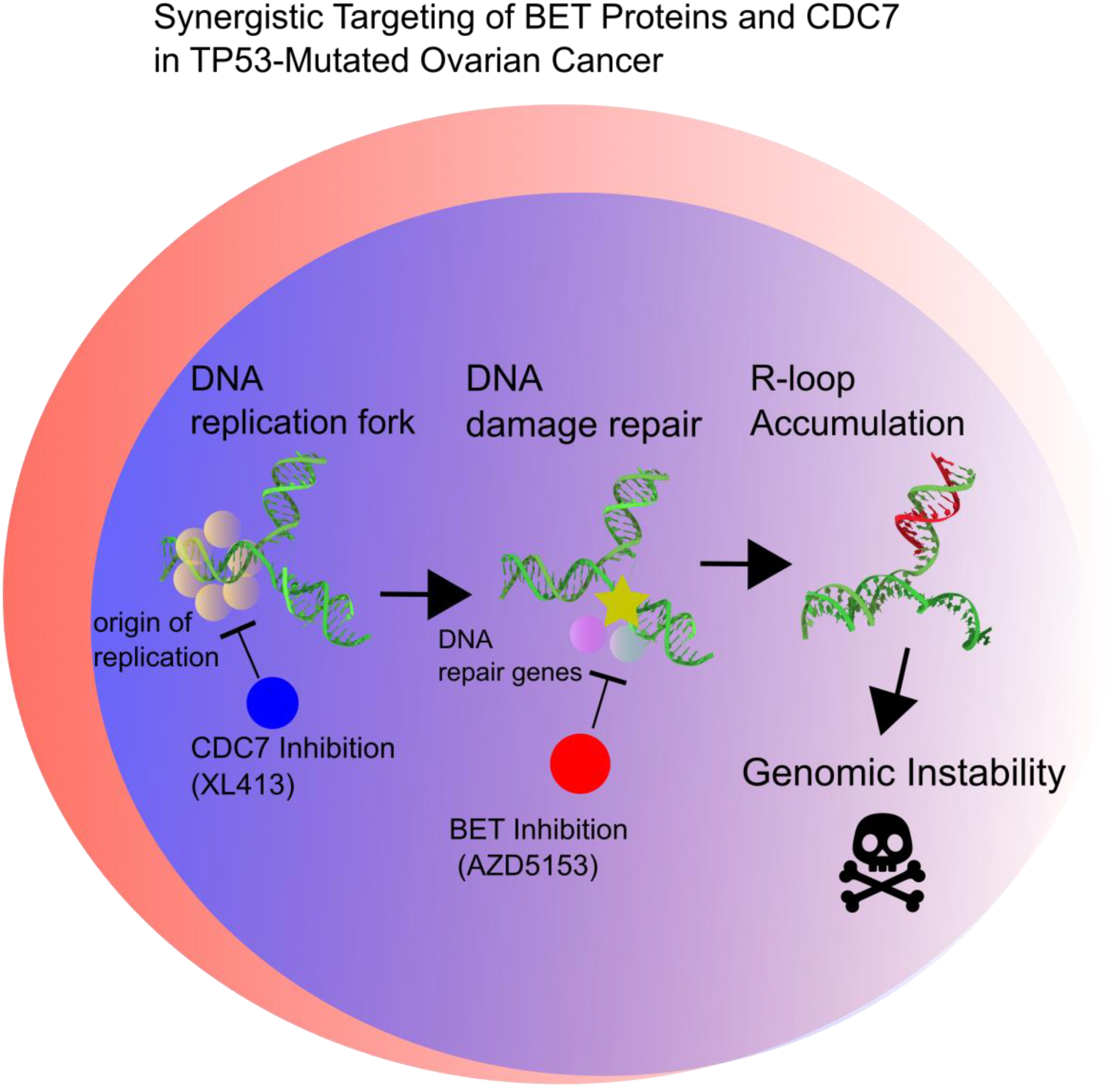

The graphic illustrates how the CDC7 inhibitor XL413 and the BET inhibitor AZD5153 work together in TP53-deficient cancer cells. First, XL413 prevents activation of the origin of replication complex, leading to DNA damage. Second, the DNA repair machinery is unable to resolve this damage, resulting in the accumulation of unresolved R-loops and genomic instability. Ultimately, this cascade culminates in cell death.

## Introduction

BET proteins are a family of transcriptional co-regulators that play a crucial role in modulating gene expression. This family includes four members: BRD2, BRD3, BRD4, and BRDT. All BET proteins feature two N-terminal conserved tandem bromodomains and a C-terminal extraterminal domain, both of which are essential for their functionality^1^. The bromodomain binds to acetylated lysine residues on histone proteins, while the extraterminal domain interacts with transcriptional activators, such as P-TEFb, allowing the BET proteins to regulate gene expression^2^.

BET proteins are implicated in various biological processes, including cell cycle regulation, DNA repair, and embryonic development^3–6^. Among the BET proteins, BRD4 stands out as a key regulator of gene expression^7^. It actively participates in chromatin remodeling and transcriptional regulation, influencing multiple genes through direct interaction or by recruiting other proteins that can promote or inhibit gene expression^8,9,10^. Given its role in regulating oncogenes like *MYC*, BRD4 has been associated with cancer development, making it an attractive therapeutic target^11^. BET-inhibitors have shown promise in treating cancer and inflammatory diseases, with the first BET-inhibitor, JQ1, demonstrating the capacity to reduce tumor growth both *in vitro* and *in vivo*^12^. Other BET-inhibitors, such as I-BET151 and CPI-0610, have also shown potential in preclinical trials as anti-tumor and anti-inflammatory agents in preclinical studies^13^.

In ovarian cancer, BET-inhibitors have been shown to decrease the growth of cancer cells, induce replication stress and increase the sensitivity to PARP inhibitors^14^. Several BET-inhibitors have been developed and evaluated in preclinical and clinical trials^15^, demonstrating promise in reducing ovarian cancer cell proliferation and preventing disease progression, while exhibiting tolerable side effects. For instance, JQ1 significantly reduced the growth of ovarian cancer cells, decreased cancer spread and improved overall survival in mouse models^16^.

Furthermore, BRD4 has been implicated in the recruitment and stabilization of replication proteins at the DNA replication fork^4,17,18^. By interacting with CDC6, PCNA and MCM7, BRD4 plays a pivotal role in the assembly and function of the replication initiation complex^4,19^.

In this study, we elucidate the intricate interplay between the BET proteins and the pre-replication complex in the context of replication stress response. By combining the BET-inhibitor AZD5153 with a reduction in origin licensing or replication initiation, we demonstrate a potent sensitization to treatment, underscoring the therapeutic potential of targeting both BET proteins and the replication processes.

## Methods

### Bioinformatics

Data from the TCGA Ovarian cancer cohort were obtained from the XENA portal (UCSC) and cBio-Portal. Expression and drug response data for cell lines were sourced from the Dependency Map portal (depmap.org). Synergies were calculated using SynergyFinder 2.0. Data analysis was performed in R, with graphical representations created using GraphPad Prism 7 and complex heatmap R packages.

### Western Blot

Cells were lysed in RIPA buffer with protease inhibitors, and protein concentration was quantified using the Bradford assay (Bio-Rad). Proteins (20 µg) were separated on a 4-15% SDS-PAGE gel, transferred to PVDF membranes (TurboBlot, Bio-Rad), and blocked with 5% BSA in TBS-Tween. Membranes were incubated with primary antibodies (Supplementary Table S1) overnight at 4°C, followed by HRP-conjugated secondary antibodies. Protein bands were detected using Clarity Chemiluminescent HRP Reagent (Bio-Rad) and visualized with a Fusion SL imaging system.

### Cell Culture

FT189, FT237, FT240, FT246, FT282, KURAMOCHI, OVCAR4, OVCAR8, and OVSAHO cells were provided by Ronny Drapkin (UPenn), while OVCAR3 and HEYA8were obtained from ATCC. Cells were cultured in DMEM F12 (Invitrogen) supplemented with 10% FBS (Atlanta Biologicals) and 1% penicillin/streptomycin (Invitrogen) at 37°C with 5% CO₂.

### Short-interfering RNA transfection

P53, MCM7, or CDT1 expression was downregulated using a pool of three siRNA duplexes (Trilencer-27, Origene), with scrambled siRNA as a negative control. Cells (1 × 10⁵/well) were transfected 24 hours post-seeding using Lipofectamine RNAiMAX (Invitrogen) with 10 nM siRNA, following the manufacturer’s protocol.

### Cell Survival Assay

Cell survival was assessed using the CellTiter 96® Non-Radioactive Cell Proliferation Assay (Promega) at 570 nm absorbance. Experiments were performed in triplicate and repeated three times. Data are presented as means ± SD, with statistical significance determined by ANOVA and post-hoc analysis. Crystal violet staining was used to count attached cells.

### Immunofluorescence

Cells on coverslips were fixed in 4% paraformaldehyde, permeabilized with 0.3% Triton X-100, and blocked. Primary antibodies were incubated for 2 hours at room temperature or overnight at 4°C, followed by secondary antibodies conjugated to Alexa Fluor dyes (Molecular Probes). Nuclei were stained with DAPI, and cells were mounted with Fluoromount-G (Sigma-Aldrich). Imaging was performed using a Zeiss Axio Observer microscope (40x). Foci and nuclear intensity were quantified using Cell Profiler and ImageJ.

### EdU Labeling

Cells were pulsed with 10 µM EdU for 20 minutes, permeabilized with 0.5% Triton X-100, and developed using a cocktail containing Tris-buffered saline, CuSO₄, Sulfo-Cyanine 3 Azide, and Sodium Ascorbate. Cells were rinsed and analyzed.

### Fiber Spread Assay

Cells were labeled with CIdU (38 µM) for 30 minutes, washed, and incubated with IdU (250 µM) for 30 minutes. Cells were lysed on slides, fixed in methanol/acetic acid, and denatured in 2.5 M HCl. Primary antibodies (anti-BrdU/IdU and anti-BrdU/CIdU) were incubated overnight, followed by secondary antibodies (Alexa 488 and Cy3). Slides were mounted with Antifade Gold (Invitrogen) and analyzed using a Leica fluorescence microscope (100x) and ImageJ.

### Cell Cycle Analysis

Cells were fixed in ice-cold ethanol, stained with DAPI/Triton X-100 solution, and analyzed using a flow cytometer (355-450 nm).

### Real-Time PCR

RNA was isolated using the RNeasy Plus Mini Kit (Qiagen), reverse-transcribed to cDNA (RT2 Easy First Strand Kit, Qiagen), and quantified using POWERUP SYBR MASTER MIX (Life Technologies) on a QuantStudio Real-Time PCR system (Thermo Fisher). Gene expression was calculated using the ΔΔCt method.

## Results

### BET-inhibitor sensitivity and origin loading

Building on previous studies that highlighted an interaction between pre-replication complex (pre-RC) proteins and BET proteins, we investigated the potential of a mutual dependency between the pre-RC and BET gene expression^4,19^. We analyzed pre-RC and BET expression data from the TCGA ovarian cancer dataset revealing that patients clustered into two main distinct groups: those with low pre-RC expression (Cluster 1) and high pre-RC expression (Cluster 2) those with (Figure 1A). We found only a weak positive correlation between the expression of the individual BET genes and origin of replication genes. To better assess this relationship, we therefore generated a combined expression score for the BET genes *BRD2*, *BRD3*, *BRD4* and *BRDT*. We found that the BRD-high tumors exhibited significantly elevated levels of *MCM2*, *MCM3*, *MCM4*, *MCM7*, and *CDC7* compared to BRD-low tumors (Figure 1B). To determine if this co-expression indicates a mutual dependency, we examined the expression of origin of replication genes and the sensitivity to a panel of various BET-inhibitors from the PRISM dataset and the Cancer Cell Line Encyclopedia (CCLE) (Figure 1C)^20,21^. Similarly, to the TCGA data, we observed a positive correlation between the expression of pre-RC genes and BET genes (Figure 1D). Interestingly, while we did not observe a strong correlation between BET gene expression and BET-inhibitor sensitivity, we noted negative correlations between the expression of pre-RC genes and sensitivity to BET-inhibitor (Figure 1D, E). This negative correlation was observed for all six BET-inhibitors analyzed, but not when assessing the sensitivity to the loss of *BRD4* via CRISPR/CAS9. This suggests that cells with high pre-RC gene levels are more sensitive towards BET-inhibitors but not to the loss of *BRD4* itself. Interestingly, a previous study demonstrated that the BET homedomain inhibitor AZD5153 does not disrupt the pre-replication complex, unlike the loss of *BRD4*^19^. Moreover, the loss of *CDC7* by CRISPR/CAS9 showed a negative correlation with sensitivity to BET inhibitors, particularly with AZD5153 (Figure 1D and F). This implies that cells sensitive to the loss of *CDC7* are more resistant to BET-inhibition and vice versa. Since the effects of BET-inhibitors can be highly cell type specific, we explored if a similar pattern could be observed in other cancer types ^22^. While we found negative correlation of pre-RC gene expression with sensitivity to BET-inhibitors in colorectal cancer, we did not find a correlation between the sensitivity to *CDC7* loss by CRISPR/CAS9 and the sensitivity to BET-inhibitors (Supp Figure 1A-D), indicating that this is a tumor type specific phenomenon. While the sensitivity towards BET-inhibitors have been linked to MYC expression in various cancer types, we did not observe such a connection in ovarian cancer cell lines in contrast to melanoma, non-small cell lung cancer or renal cell carcinoma (Figure 1D, Supp. Figure 1D, E)^12^.

**Figure 1:**
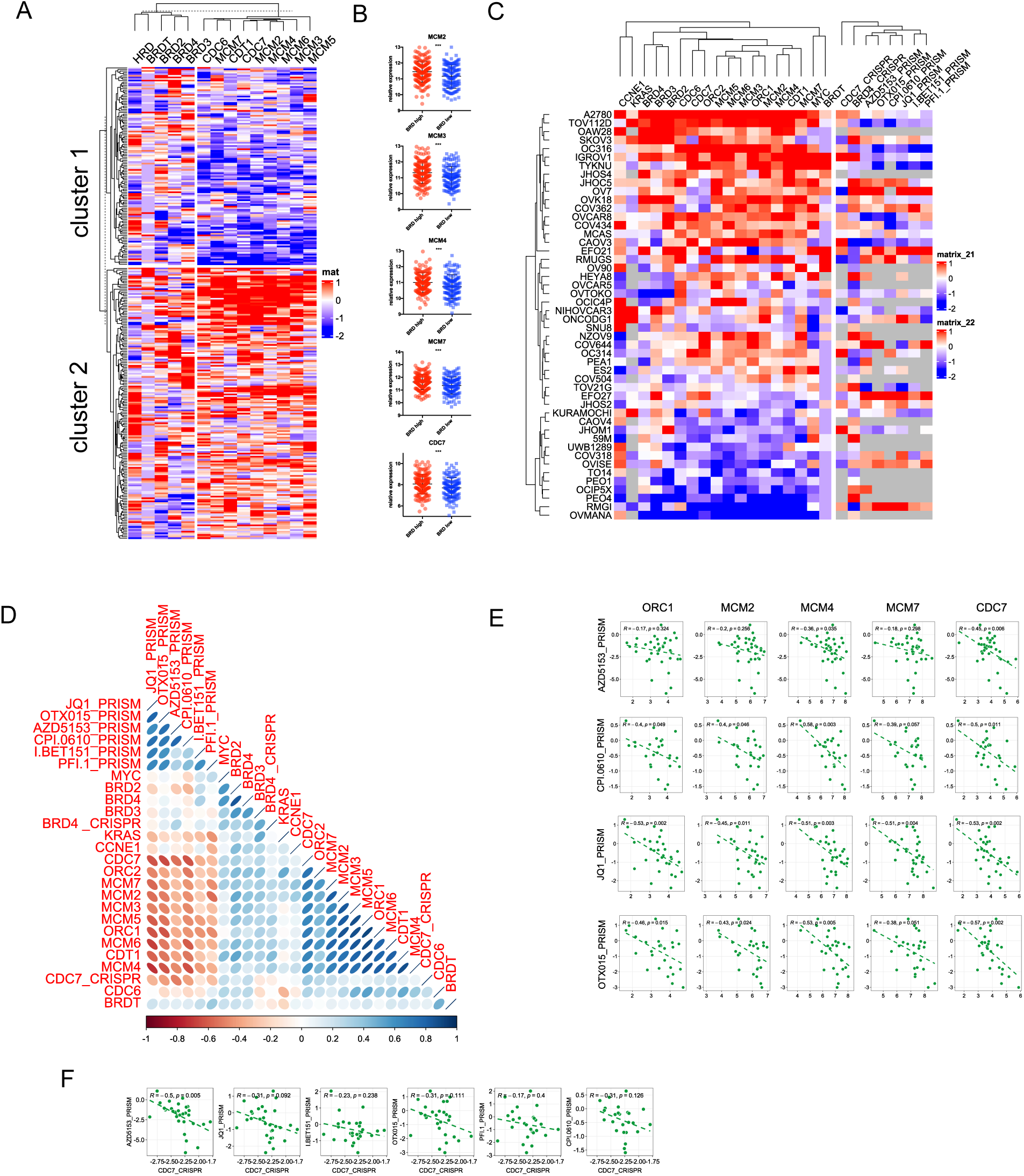
BET-inhibitor sensitivity and origin loading. A) Heatmap analysis of BRD genes and origin of replication genes in the ovarian cancer dataset of the TCGA data. B) Expression of origin of replication genes in patients expressing high or low levels of BRD genes. C) Ovarian cancer cell line gene expression from the CCLE dataset and dependency scores of different BRD inhibitors and CRISPR dependency score from the PRISM dataset. A low dependency score indicates a higher sensitivity. D) Correlation matrix from the data from C. E) Individual values of ORC1, MCM2, MCM4, MCM7 and CDC7 expression against the PRISM dependency score of AZD5153, CPI.0610, JQ1 and OTX015. Depicted are pearson correlations with the corresponding p-values. F) Individual values of the CDC7 CRISPR dependency score against the PRISM dependency score of AZD5153, JQ1, I.BET151, OTX015, PFI.1 and CPI.0610. Depicted are pearson correlations with the corresponding p-values.

In summary, our results indicate a mutual dependency between the expression of pre-RC genes and the sensitivity towards BET-inhibitors in ovarian cancer.

### Synergistic effects of AZD5153 with XL413

Based on our findings, indicating a link between the pre-RC expression and the sensitivity to BET-inhibitors, we investigated whether knocking down pre-RC genes by siRNA would alter the sensitivity of ovarian cancer cells to BRD4 inhibitors.

We utilized the BET-inhibitor AZD5153 (BRD4i) alongside siRNA targeting either *MCM7* or the licensing factor *CDT1*, both of which, along with *CDC6* play critical roles in the formation of the pre-replication complex. Our results showed a significant sensitization to AZD5153 in cells treated with *CDT1* siRNA, while *MCM7* siRNA treatment resulted in a more modest sensitization (Figure 2A, B).

**Figure 2:**
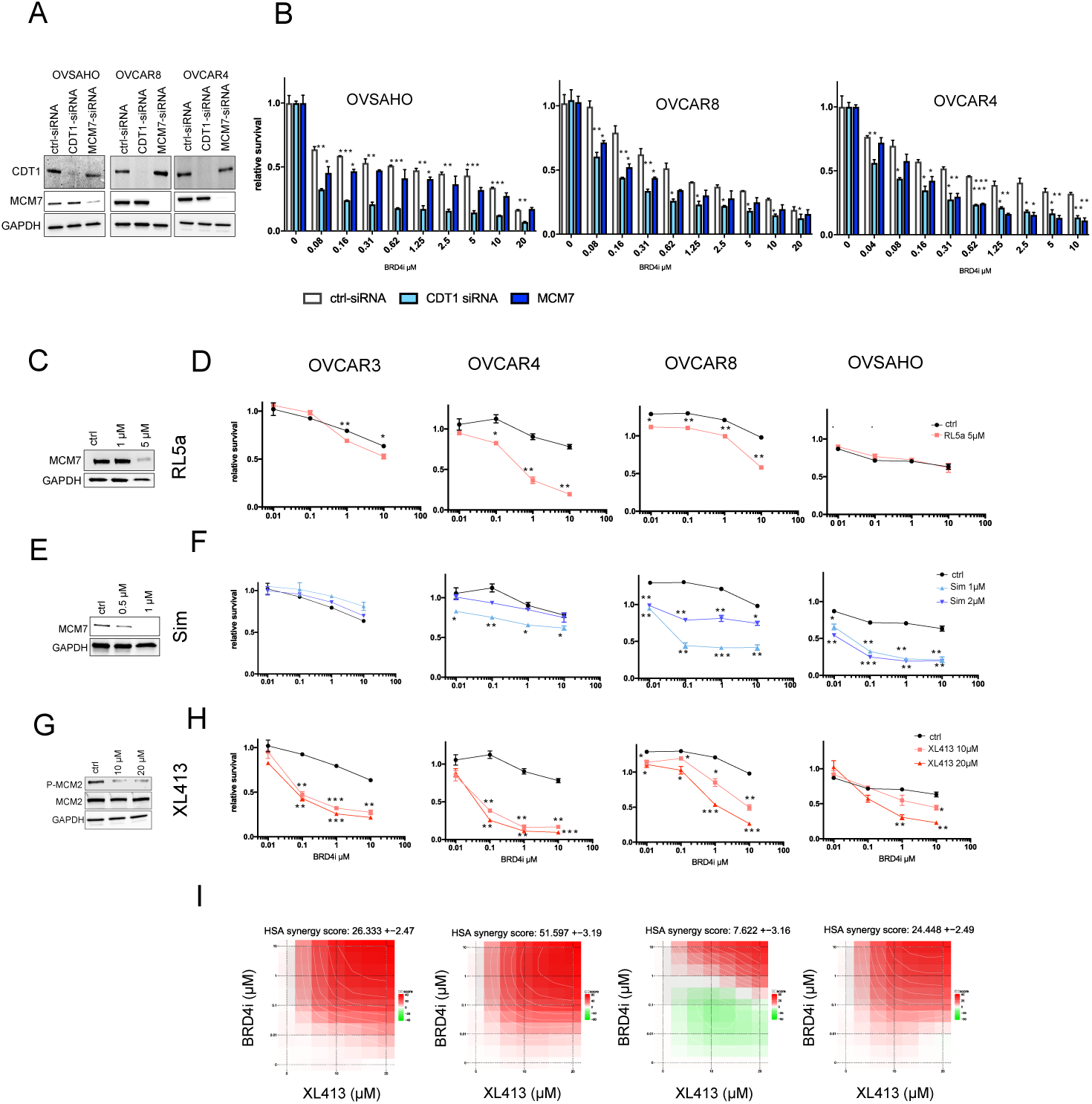
Synergistic effects of AZD5153 with RL5a, Simvastatin and XL413. A) Western blot analysis of OVSAHO, OVCAR8 and OVCAR4 cells treated with control siRNA or siRNA targeting CDT1 or MCM7. Blots were probed against CDT1, MCM7 and GAPDH. B) Relative survival in cells described in a) and treated with different concentrations of the BRD4i AZD5153 for 96 hours. Representative western blot analysis of OVCAR8 cells treated with control or RL5a (1µM and 5µM) for 72 hours. Blots were probed against MCM7 and GAPDH. D) Relative survival in cells described in C) and pre-treated with 5µM RL5a. After 72 hours treatment with RL5a the cells were treated with different concentrations of the BRD4i AZD5153 for 96 hours before survival was measured. E) Representative western blot analysis of OVCAR8 cells treated with control or simvastatin (0.5µM and 1µM) for 72 hours. Blots were probed against MCM7 and GAPDH. F) Relative survival in cells pre-treated as described in E). After 72 hours treatment with simvastatin cells were treated with different concentrations of the BRD4i AZD5153 for 72 hours before survival was measured. G) Representative western blot analysis of OVCAR8 cells treated with control or XL413 (10µM and 20µM) for 72 hours. Blots were probed against P-MCM2, MCM2 and GAPDH. H) Relative survival in cells pre-treated as described in G. After 72 hours treatment with XL413 cells were treated with different concentrations of the BRD4i AZD5153 for 72 hours before survival was measured. I) Depicted are synergy blots with calculated HSA synergy scores from the samples shown in H).

To further explore the potential of modulating replication origins and initiation as a strategy to enhance the efficacy of AZD5153, we conducted pre-treatment experiments using drugs that impact pre-replication proteins or replication initiation. Specifically, we used RL5a, a compound known to reduce the expression of the pre-RC genes *MCM2-7*^23^. Following a 72-hour pre-treatment with RL5a, we observed increased sensitivity to AZD5153 in OVCAR4 and OVCAR8 cells, while no sensitization was observed in OVSAHO and OVCAR3 cells (Figure 2C, D).

Additionally, we examined the effects of statins, which have been reported to decrease MCM protein levels^24,25^. Pre-treatment with simvastatin sensitized OVCAR4, OVCAR8, and OVSAHO cells to AZD5153; however, the synergistic effect was only observed within a narrow concentration window of simvastatin. Notably, some cell lines, such as OVCAR8, exhibited high sensitivity to simvastatin on its own (Figure 2E, F). In contrast, the CDC7 inhibitor XL413, which disrupts the phosphorylation of MCM2 and its replication origin activation, demonstrated a robust synergistic effect across a wide range of concentrations in all tested cell lines (Figure 2G-I)^26^.

### The synergistic effects are partially dependent on TP53 mutation

Next, we examined whether this synergistic effect could also be observed in immortalized fallopian tube cell lines^27^. To our surprise, all four tested cell lines exhibited strong synergy when pre-treated with various concentrations of the CDC7 inhibitor XL413, followed by the addition of the BET inhibitor AZD5153 (Figure 3A). Importantly, these fallopian tube cell lines were immortalized by inhibiting the *TP53* gene either through the expression of the SV40 Large T antigen (in FT189) or via shRNA-mediated knockdown of P53 (in FT237, FT240, and FT246). To determine if the synergistic effects depend on the disruption of the *TP53* pathway, we utilized a panel of primary ovarian cancer cell lines. These lines either contained accumulated P53 (indicating a *TP53* mutation -MA1-MA6 and MA11) or expressed normal P53 levels (MA7, MA8, MA9, MA10 and MA12) (Figure 3B). We found that the synergistic effect of XL413 and AZD5153 was observed only in the cell lines with accumulated P53, which is indicative of mutated *TP53* (Figure 3B, C). Since the primary cell lines that were not affected by the combinational treatment also lacked PAX8 expression, a marker for high-grade serous carcinomas (HGSC), we could not rule out the possibility that the observed effect might be subtype-specific for reasons other than TP53 disruption (Figure 3B). To further investigate this, we used HeyA8 cells, which express wild type P53 and only show a weak sensitization to AZD5153 when treated with XL413 (Figure 3D). Upon P53 knockdown using siRNA we observed a marked enhancement of the synergistic effect between XL413 and AZD5153, resulting in a 100-fold increase in sensitivity to AZD5153. This indicates that the synergy is indeed dependent on the loss of *TP53* (Figure 3D).

**Figure 3:**
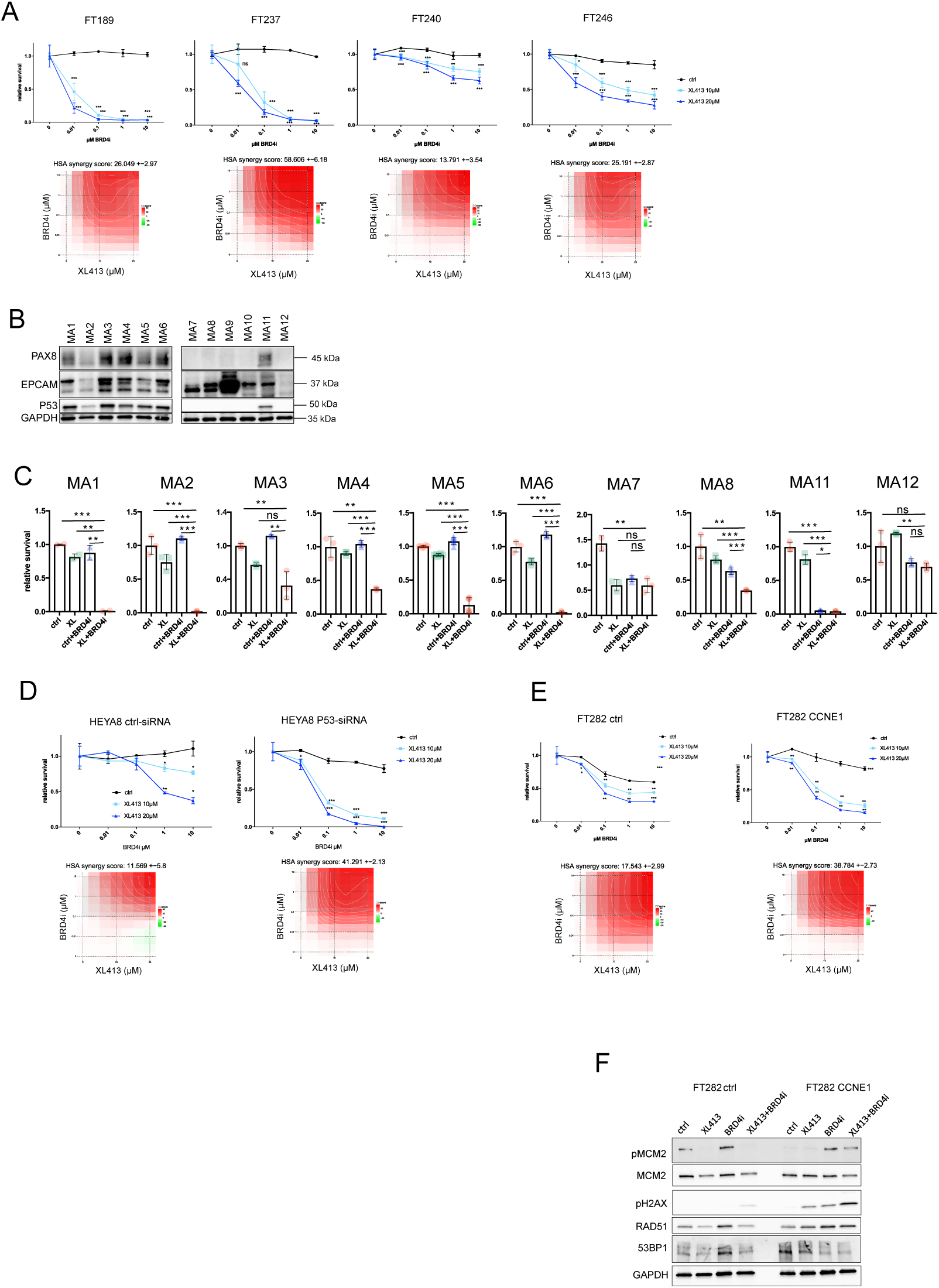
The synergistic effects are partially dependent on TP53 mutation. A) Upper panel: Relative survival of FT189, FT237, FT240 and FT246 cells after pre-treatment with 10µM or 20µM XL413 for 72 hours before adding different concentrations of the BRD4i AZD5153. Lower panel: HSA synergy analysis of corresponding cell lines in the upper panel. B) Western blot analysis of a panel of primary cell lines and analyzed for PAX8, EPCAM, P53 and GAPDH. C) Relative survival of primary cells from b) pre-treated with 10µM XL413, followed by the treatment of 0.1µM AZD5153 for each 72 hours. D) Upper panel: Relative survival of HEYA8 cells that were treated with control or siRNA against P53 after pre-treatment with 10µM or 20µM XL413 for 72 hours before adding different concentrations of the BRD4i AZD5153. Lower panel: HSA synergy analysis of corresponding cell lines in the upper panel. E) Upper panel: Relative survival of FT282 cells that express normal or high levels of Cyclin E1 (CCNE1) after pre-treatment with 10µM or 20µM XL413 for 72 hours before adding different concentrations of the BRD4i AZD5153. Lower panel: HSA synergy analysis of corresponding cell lines in the upper panel. F) Western blot analysis of samples analyzed in E) for expression of p-MCM2, MCM2 pH2AX, RAD51, 53BP1 and GAPDH.

We also explored if the expression of other genes such as cyclin E1, (*CCNE1*) influences the synergistic effect. *CCNE1* is frequently amplified in HGSCs (20%) and its overexpression has been shown to disrupt the origin loading^28–30^. We employed the immortalized fallopian tube cell line FT282, which expresses shRNA against *TP53* as well as an inducible *CCNE1* vector. While we observed a synergistic effect in cells with normal cyclin E1 levels, this synergy was significantly enhanced in cells overexpressing cyclin E1 (Figure 3E).

Lastly, when analyzing the effects on DNA damage signaling, we observed a strong increase phosphorylated yH2AX (p-yH2AX) in cyclin E1 overexpressing cells following double treatment, while levels of 53BP1 decreased (Figure 3F).

These findings suggest that the synergistic effects of XL413 and AZD5153 are partially dependent on TP53 status and may be influenced by the expression of genes involved in cell cycle regulation, such as cyclin E1.

### Effects on DNA damage signaling pathways

To assess the impact on DNA damage signaling pathways, we examined the effects of MCM7 knock down and AZD5153 on various markers of DNA damage. Remarkably, treatment with MCM7 siRNA in conjunction with AZD5153, resulted in a significant increase in p-yH2AX levels, a key indicator of DNA damage. Additionally, we observed a decrease in nuclear 53BP1 foci staining, which represents a marker of DNA double-strand break (DSB) repair (Figure 4A) Intriguingly, MCM7 knockdown also led to a substantial increase in RAD51 foci, but this was abolished upon treatment with the BET-inhibitor. Next we tested if the combination of AZD5153 and the CDC7 inhibitor XL413 had a similar effect on DNA damage markers. In contrast to RNAi treatment, we did not observe an additive or synergistic increase in p-yH2AX with the combination treatment (Figure 4B). Similarly to the effects seen with *MCM7* RNAi, the number of 53BP1 foci decreased when comparing the combination of AZD5153 with XL413 to XL413 treatment alone (Figure 4C).

**Figure 4:**
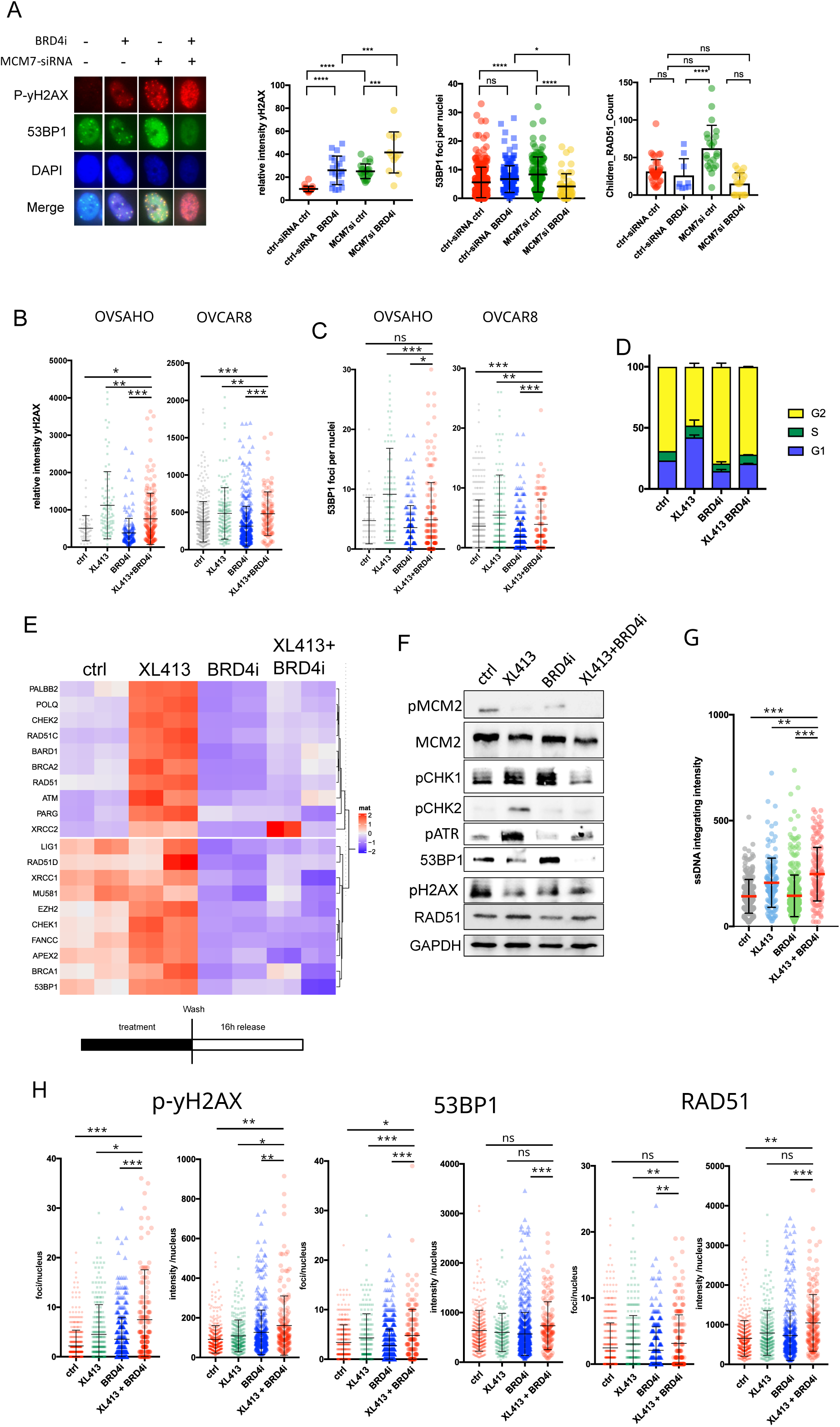
Effects on DNA damage signaling pathways. A) Immunofluorescence of OVCAR8 cells treated with MCM7-siRNA followed by BRD4i treatment. Cells were analyzed and quantified for p-yH2AX (red), 53BP1 (green) and nuclear DNA (blue). RAD51 foci were additionally stained, analyzed and counted. B) p-yH2AX Immunofluorescence analysis of cells treated with XL413 followed by BRD4i treatment. C) 53BP1 Immunofluorescence analysis of cells treated with XL413 followed by BRD4i treatment. D) Cell cycle analysis of cells treated with XL413 followed by BRD4i. E) Heatmap of real time PCR analysis of cells treated with XL413 followed by BRD4i. F) Western blot analysis of OVCAR8 cells treated with XL413 followed by BRD4i. Blot was analyzed for pMCM2, MCM2, pCHK1, pCHK2, pATR, p-yH2AX, RAD51, and GAPDH. G) relative nuclear fluorescence intensity of single stranded DNA (cIdU) of OVCAR8 cells treated with XL413 followed by BRD4i treatment. H) Upper scheme: After treatment regime, cells are washed and released for 16 hours into normal media before immunofluorescence was performed. Lower panel: Analysis of foci per nuclei and integrated intensity per nuclei of p-yH2AX, 53BP1 and RAD51 in OVCAR8 cells.

We also analyzed effects on the cell cycle and found that XL413 resulted in a strong accumulation of cells in the G1 phase due to G1 checkpoint arrest. However, this accumulation was diminished by AZD5153 treatment (Figure 4D). We next performed real-time PCR using a gene panel to determine whether other DNA damage-related genes were affected by the treatment of AZD5153 and XL413 (Figure 4E). Recent studies have shown BET-inhibitors can reduce the gene expression of several genes involved in DSB repair^14^. Our real-time PCR analysis confirmed that XL413 treatment led to the upregulation of several genes involved in double-strand break repair, including *RAD51*, *53BP1*, *RAD51D*, *POLQ*, *BARD1*, *RAD51C*, *BRCA1*, *BRCA2*, and *XRCC2* (Figure 4E). Notably, the upregulation was reversed by the addition of AZD5153. This finding aligns with our observation that while XL413 activates the serine/threonine kinase CHK1, CHK2 and ATR, this activation was abrogated by the combination with AZD5153 (Figure 4F).

To further explore replication stress in cells treated with both drugs, we performed native BrdU immunofluorescence to measure the levels of single-strand DNA (ssDNA), which serves as a marker for replication stress. Importantly, cells treated with both XL413 and AZD5153 exhibited significantly higher levels of ssDNA compared to cells treated with either agent alone or control cells (Figure 4G).

To assess if the lack of DNA damage response and continuous progression through the cell cycle leads to the accumulation of more unresolved DNA damage during the combination treatment, allowing the cells to recover in normal media for 16 hours after the standard treatment protocol (Figure 4H).

Remarkably, upon recovery, we noted a pronounced elevation in p-yH2AX, 53BP1 and RAD51, indicating that cells accumulate more DNA damage during the combination treatment.

These findings suggest that while XL413 treatment alone enhances DNA damage and DNA damage response, combining XL413 with AZD5153 exacerbates genomic instability by impairing the cells’ ability to effectively respond to DNA damage and replication stress.

### Reduction in origin licensing and BRD4 inhibition leads to an increase in R-loops

Recent studies have shown that BET-inhibition can lead to an increase in R-loop formation, which refers to DNA:RNA hybrids to occur as a result of conflicts between the transcription and the replication machinery^17,31^. Building on this knowledge, we aimed to investigate whether combinating BET-inhibition with a reduction in origin licensing or replication origins could enhance R-loop accumulation, thus elucidating the underlying mechanism behind the observed increase in replication stress.

Indeed, when treating cells with MCM7 siRNA in combination with AZD5153, we observed a significant increase in nuclear S9.6 staining, an antibody that specifically binds to RNA:DNA duplexes (Figure 5A). This effect was similarly observed across multiple cell lines when XL413 was combined with AZD5153 (Figure 5B). In contrast to findings in other studies, we noted that treatment with the BET inhibitor alone resulted in only a weak or negligible increase in R-loop formation in our analyzed cell lines^31^. Similarly, to the XL413 treatment, we also observed an increase in R-loop formation when combining RL5a with the AZD5153 (Sup-Figure 2A). We next investigated whether cyclin E1 levels influenced R-loop formation.

**Figure 5:**
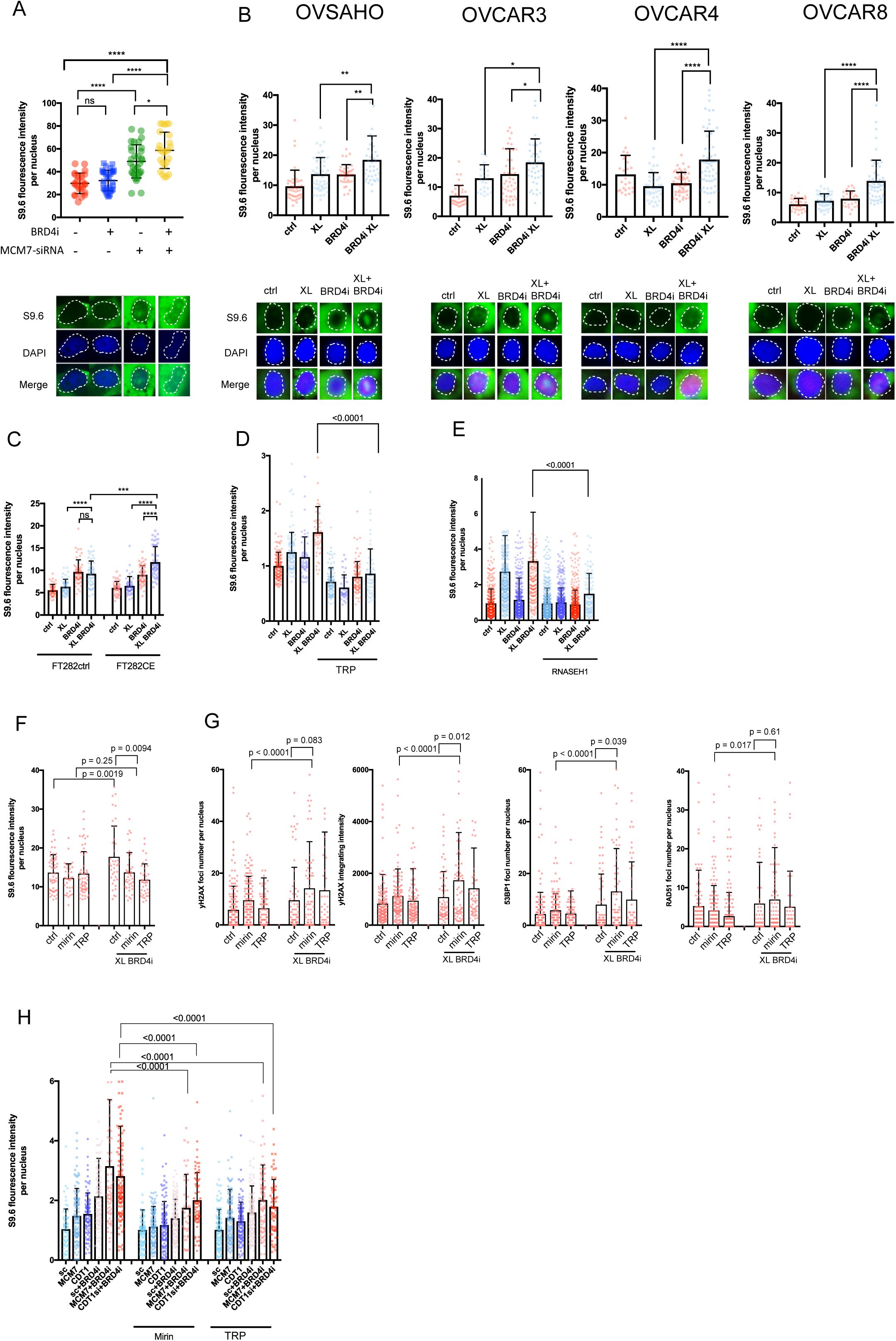
Reduction in origin licensing and BRD4 inhibition leads to an increase in R-loops. A) Nuclear S9.6 immunofluorescence analysis of OVCAR8 cells treated with control or MCM7-siRNA followed by BRD4i treatment. B) Nuclear S9.6 immunofluorescence analysis of different cell lines treated with XL413 followed by BRD4i treatment. C) Nuclear S9.6 immunofluorescence analysis of FT282-ctrl and FT282-CCNE1 cells treated with XL413 followed by BRD4i treatment. C) Nuclear S9.6 immunofluorescence analysis of OVCAR8 cells treated with XL413 followed by BRD4i treatment. Prior to BRD4i treatment, cells were treated with the TFIIH inhibitor Triptolide (TRP) to block transcription initiation. E) Nuclear S9.6 immunofluorescence analysis of OVCAR8 cells treated with XL413 followed by BRD4i treatment. Prior to treatment, cells were transduced with a control vector or a vector expressing RNASEH1. F) Nuclear S9.6 immunofluorescence analysis of OVCAR8 cells treated with XL413 followed by BRD4i treatment. Prior to BRD4i treatment, cells were treated with Mirin or TRP. G) Immunofluorescence analysis of OVCAR8 cells treated with XL413 followed by BRD4i treatment. Prior to BRD4i treatment, cells were treated with Mirin or TRP for 48 hours before cells were washed and released into normal media for 16 hours. Cells were analyzed for p-yH2AX foci number and integrated nuclear intensity, as well as 53BP1 and RAD51 foci numbers per nucleus. H) S9.6 immunofluorescence analysis of OVCAR8 cells treated with MCM7 or CDT1-siRNA followed by BRD4i treatment. Prior to BRD4i treatment, cells were treated with TRP or Mirin.

Using immortalized fallopian tube cell line FT282, we found that in cells expressing low levels of cyclin E1, there was no increase in R-loops upon combined treatment. In contrast, cells with high cyclin E1 expression exhibited a significant rise in R-loop levels (Figure 5C). To assess the transcription dependence of this effect, we employed Triptolide (TRP), a TFIIH inhibitor that blocks transcription initiation and leads to RNA polymerase II (*RNAPII*) degradation. Interestingly, TRP treatment reduced S9.6 nuclear fluorescence staining in cells treated with both XL413 and AZD5153 (Figure 5D). Moreover, overexpressing RNase H1, an enzyme responsible for removing R-loops, resulted in decreased R-loop formation (Figure 5E).

Recently a dual role of the MRN (Mre11, Rad50 and Nbs1) complex in R-loop formation has been suggested. On one hand, it can resolve R-loops present at the replication fork through a nucleolytic-independent mechanism. On the other hand, it can promote R-loops to facilitate the resection at DSBs and thereby enhances homologous recombination in conjunction with RNAPII^32–34^. To investigate if the increase in R-loop formation could be attributed to enhanced DNA end-resection, we pretreated the cells with the MRE11 inhibitor Mirin. Encouragingly, Mirin treatment reduced S9.6 nuclear intensity in cells subjected to the combination of AZD5153 with XL413 (Figure 5F). These results support the model that both, MRN and RNAPII, are required for R-loop induced double strand break^33^. They also align with findings that resolution of R-loops is facilitated by the homologous recombination and Fanconi armenia genes, both of which are downregulated in our model following AZD5153 treatment^35^. This is further corroborated by the research from Ohle et al, which indicates that R-loops are essential efficient HR-dependent double strand repair. We then tested whether inhibiting R-loops and the MRN complex would reduce the ability of cells to repair DSBs by HR^33^. We therefore treated the cells additionally to our combinational treatment with the MRN inhibitor Mirin or TRP before allowing the cells to recover in normal media for 16 hours. Following Mirin treatment, we observed a significant increase in p-yH2AX foci number and nuclei intensity (Figure 5G). When analyzing the number of RAD51 and 53BP1 foci we observed that upon the treatment with mirin the cells increased 53BP1 foci numbers whereas the number RAD51 foci remained unchanged. This indicates a shift from HR DSB repair to alternative DSB repair pathways that require no DNA end-resection.

To confirm our findings, we examined if inhibiting the MRN complex or RNAPII could also decrease nuclear S9.6 in cells treated with *MCM7* or *CDT1* siRNA alongside the BRD4 inhibitor AZD5153. Indeed, inhibiting both the MRN complex or RNAPII resulted in a significant reduction in R-loop formation in cells co-treated with AZD5153 (Figure 5H). Collectively, these data demonstrate that the combined treatment of XL413 and AZD5153 promotes an increase in genomic instability that is caused by unresolved DSBs. The MRN complex together with RNAPII promotes end resection and R-loop formation that can’t be resolved in the double treated cells.

### Reduction in origin licensing and BRD4 inhibition effects replication fork stability

Based on the previous results we investigated the effects of combined treatment with XL413 with AZD5153 on replication fork stability using a fiber combing assay (Figure 6A). The combined treatment significantly reduced the number of ongoing replication forks by increasing fork stalling (Figure 6B). Notably, after the inhibitors were removed, there was an increase in origin firing. We also observed a significant increase in replication speed, further supporting our finding that R-loops are increased DSBs rather than at the replication fork, since previous research indicated that R-loops at the replication fork slow down replication^32^ (Figure 6C).

**Figure 6:**
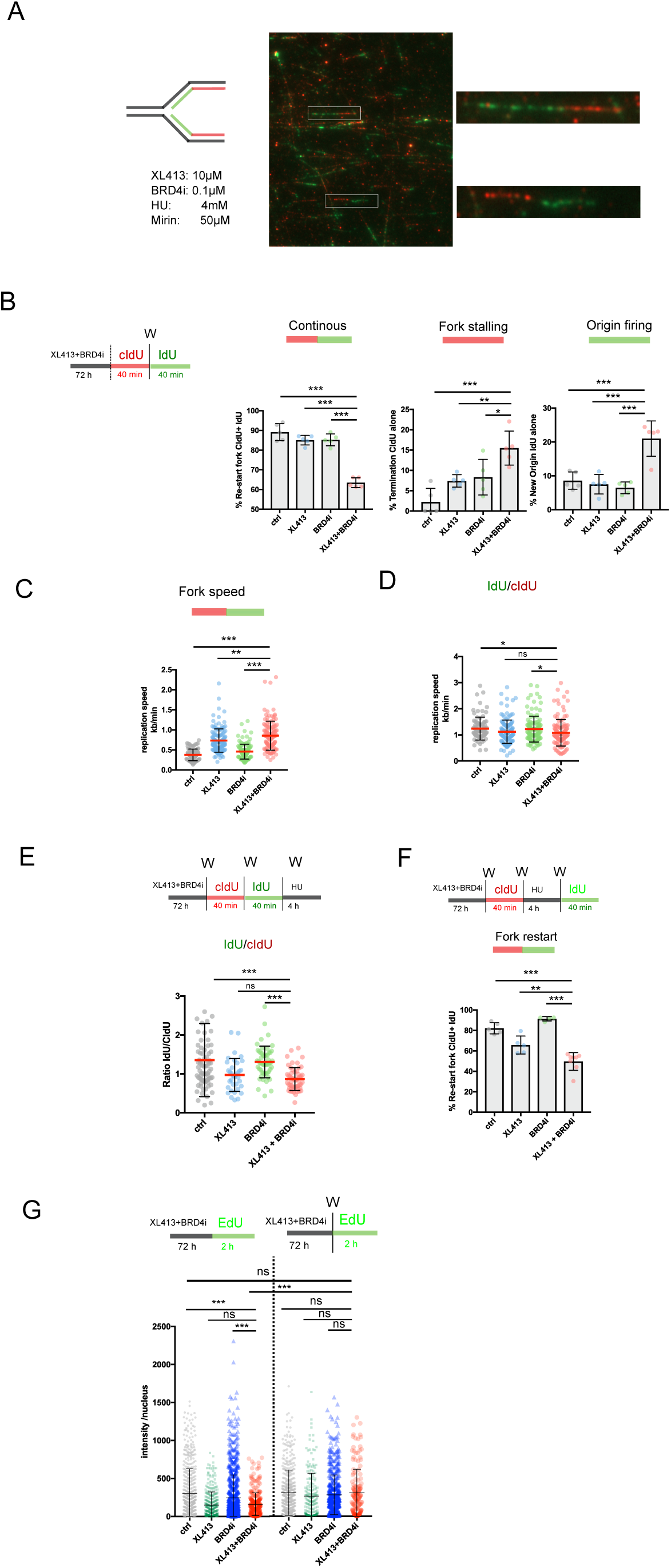
Reduction in origin licensing and BRD4 inhibition affects replication fork stability. A) Left: Sketch of a replication fork treated with cIdU (red) followed by IdU (green). Below are the concentrations of the used substances in this figure. Right: A representative immunofluorescence image of a fiber spread assay with magnified fibers displayed. B) Left: treatment order of XL413 and BRD4i followed by cIdU for 40 minutes, wash steps (W) and 40 minutes of IdU treatment. Right: analyzed fibers for continuous forks, stalling forks and new origin firing. C and D) Fibers from B) were analyzed replication speed in kb/min C) and fork stability was analyzed by calculating the ratio of IdU to cIdU for each fiber D). E) Upper panel: treatment order. Lower panel: fork stability was analyzed by calculating the ratio of IdU to cIdU for each fiber. F) Upper panel: treatment order. Lower panel: fork restart was analyzed. G) Upper panel: treatment order. On the left, EdU was directly added to the cells whereas on the right cells were washed (W) before adding EdU. Lower panel: quantified EdU intensity per nucleus.

Increased replication speed is associated with genomic instability and can occur due to either the upregulation of proteins that enhance transcription or a compensatory mechanism where increased replication speed offsets the greater distance between replication origins^36^. In contrast, fork stability was only weakly affected by the combined treatment (Figure 6D). The increase in replication speed could also be attributed partially to ATR activation following XL413 treatment when compared to control (Figure 4F).

For further analysis of fork stability, we treated cells sequentially with cIdU and IdU for 30 minutes each, followed by a high dose of 4mM hydroxyurea (HU) for 4 hours. Cells exhibiting reduced fork stability will display a lower IdU to cIdU ratio. While we observed a decreased fork stability with both XL413 alone and in combination with AZD5153, there was no significant difference between the two treatments (Figure 6E), suggesting that XL413 is the primary contributor to replication fork instability.

We next examined the ability of cells to restart replication after HU treatment. Cells subjected to the combined treatment with XL413 and AZD5153 were less capable of restarting replication (Figure 6F).

During our analysis of cells in S-phase, using the EdU incorporation assay, we found that both XL413 and the combined treatment significantly reduced the number of replicating cells (Figure 6G). However, by washing out the treatment before administering EdU, this effect was mitigated (Figure 6G). In summary, the combined treatment of XL413 and AZD5153 results in increased fork stalling and enhanced replication speed, highlighting changes in the intricate interplay of fork dynamics and genomic instability.

## Discussion

The bromodomain and extraterminal (BET) protein family, consisting of BRD2, BRD3, BRD4, and BRDT, functions as epigenetic readers that specifically recognize acetylated lysine residues on chromatin^37^. Among these proteins, BRD4 is the most extensively studied with established roles in multiple cancer types, including ovarian cancer^38–41^. Its primary role in cancer involves regulating transcription through the recruitment of critical regulatory elements^39^. Specific inhibitors which disrupt the interaction between BET proteins and chromatin have shown promising results in preclinical models^37,41^.

BRD4 is intricately involved in the DNA replication stress response and regulates the pre-replication complex, which is crucial for proper intra-S phase checkpoint signaling^19^. BRD4 has been shown to interact with several components of the pre-replication complex, including MCM5, MCM7, CDC6 and CDC7.

In our study, we observed a positive correlation between BET expression and origin of replication gene expression. Moreover, analysis of the Cancer Cell Line Encyclopedia dataset, indicated that increased expression of origin of replication genes exhibited greater sensitivity to BET inhibitors, suggesting a dependency on these genes. Interestingly, this correlation was not apparent in BRD4 knockout experiments using CRISPR/CAS9, which aligns with previous research showing that the BET-inhibitor AZD5153 does not disrupt the pre-replication complex like BRD4 knockout does^19^. This in turn could contribute significantly to the sensitivity and resistance to BET inhibition. We therefore speculated that reducing the number of replication origins might sensitize cells to BET inhibitors. Indeed, our results confirmed this hypothesis: pre-treatment with siRNA targeting MCM7 or CDT1 as well as small molecules like RL5a and XL413 significantly increased sensitivity to AZD5153. Notably, the strongest synergy was observed with the CDC7 inhibitor XL413, which prevents activation of the pre-replication complex. Additionally, we found that loss of *CDC7* via CRISPR/CAS9 correlates negatively with the sensitivity towards different BET-inhibitors, particularly AZD5153.

Importantly, this synergy was markedly enhanced in cells with an inactive *TP53* pathway or elevated levels of cyclin E1, indicating a cancer cell-specific effect.

Reducing the number of replication origins increases replication stress resulting in an increased prevalence of cells in G1 phase. Echoing findings of Zhang et al. we observed a strong reduction of DNA damage signaling following the additional treatment with a BET-inhibitor both on the transcriptional and protein levels^19^. Their study demonstrated that BET inhibition reduces CHK1 activity, compromising the G2/M checkpoint and subsequently resulting in the accumulation of DNA damage^19^. This could explain the pronounced synergistic effects observed in TP53 deficient cells, as the absence of a functional G1/S checkpoint necessitates reliance on an intact G2/M checkpoint to maintain genomic integrity.

It is important to note that HGSCs commonly harbor *TP53* mutations, which are thought to be among the earliest in cancer evolution^42^. These mutations are already detected in benign precursor lesions like P53 signatures and in malignant lesions such as serous tubal intraepithelial carcinomas (STIC)^43^. Mathematical modeling indicates that it can take decades from the emergence of a *TP53* mutation to evolve into a STIC and subsequently several years to progress into a HGSC^44^. The early detection of STIC lesions or early stages of HGSC remains one of the biggest challenges. If this could be reliably achieved it would become of interest if combination therapies like the one tested here could effectively eradicate both precursor lesions as well as advanced HGSCs.

Furthermore, we found that elevated cyclin E1 levels further increased sensitivity to the combined treatment with the CDC7 inhibitor XL413 and the BETi AZD5153 even further.

In normal cells the expression of cyclins, including cyclin E1, is tightly regulated during the cell cycle. Recent studies indicate that the origin of the replication gene ORC1 represses the expression of *CCNE1* transcriptionally, while CDC6 increases it^45^. Additionally, during the G1 phase, a CDC6 motif cooperates with cyclin E1-CDK2 to promote ORC1-CDC6 interactions^46^. Beyond direct interactions with the origin of replication complexes, aberrant expression of cyclin E1 might also influence the origin of replication indirectly by shortening G1 phase limiting the cellular capacity to establish the required number of replication origins for a successful replication during S phase.

Importantly we observed a pronounced synergistic effect in immortalized fallopian tube cells expressing high cyclin E1 levels, which was accompanied by increased R-loop formation and enhanced p-yH2AX staining. These findings suggest that the observed synergy may be specific to cancer cells. Nevertheless, synergy was identified in various tested cell lines, including those with cyclin E1 amplification (OVCAR3), gain (OVCAR4), or diploid *CCNE1* status (OVCAR8, OVSAHO), indicating that other factors likely contribute to the observed synergistic effects.

Moreover, ours and previous results indicate that reducing replication initiation by the CDC7 inhibitor XL413 limits replication under BET-inhibitor treatment, implying that BET inhibition leads to aberrant DNA replication through a mechanism dependent on CDC6-CDC7^19^. In our present study we expanded upon this concept and discovered that XL413 and AZD5153 work synergistically through distinct mechanisms of action. While the treatment with XL413 induces DNA damage and upregulates DNA repair genes, AZD5153 counteracts these effects by transcriptionally downregulating many of these DNA repair genes. Consequently, the inability of cells to repair the extensive levels of DNA damage and ultimately cell death.

Another mechanism influenced by the combination treatment with XL413 and AZD5153 is the accumulation of unresolved R-loops. Recent studies have indicated that inhibition of BRD4 results in R-loop accumulation^17,31^, which manifests as the collision between the replication and transcription machinery, ultimately leading to replication stress and DNA damage. In our analysis of cancer cell lines, we observed that reducing *MCM7* expression through siRNA, as well as employing small molecules such as RL5a and XL413 in conjunction with the BET-inhibitor AZD5153 significantly increased R-loop formation. Notably, in contrast to Edwards et al, we detected only a weak or negligible increase in R-loop formation following single agent treatment with the BET-inhibitor^31^.

Utilizing the TFIIH inhibitor Triptolide (TRP), which blocks transcription initiation and leads to RNAPII degradation, we observed a corresponding reduction in R-loop formation. Overexpressing RNaseH, an enzyme that degrades the RNA moiety in R-loops, similarly reduces R-loop levels. Recent studies have highlighted the dual roles for the MRN complex in R-loop formation; it can both resolve R-loops at the replication fork through a nuclease-independent mechanism and promotes R-loops to facilitate resection at double-strand breaks (DSBs), thus aiding homologous recombination (HR) in collaboration with RNAPII^32–34^.

Our results support the model that MRN nuclease activity is crucial for promoting R-loops that regulate resection at DSBs^32,33,47^. Interestingly, the C-terminal binding protein interacting protein (CtIP), known to activate the MRN nuclease activity, has also been shown to suppress R-loop formation at sites of radiation damage^48^. It was further shown that inhibition of BRD4 induces HR-deficiency through the depletion of CtIP^14^, indicating that the loss of CtIP could play a role in R-loop formation following BET-inhibitor treatment in our model.

Overall, the combination of molecules that target CDC7 with BET-inhibitors suggests a potential strategy that might help address cancer cells through what appears to be a synthetic lethal interaction with BET-inhibitors. While the preliminary results are encouraging, further research is needed to validate these findings across different cancer types and in more clinically relevant models.

## Acknowledgements

We thank members of the Marme/Doberstein lab and the Department of Gynecology Mannheim for their valuable discussions and feedback. We are particularly grateful to Stefanie Gaiser from the Marme/Doberstein lab for her technical assistance. This work was supported by the AstraZeneca iEvsion Grant.

## Data Availability

The data supporting the findings of this study are available within the article and its supplementary materials. Additional datasets generated and/or analyzed during the current study are available from the corresponding author upon reasonable request.

## Ethics Statement

This study did not involve animal subjects. However, human samples were obtained from patients undergoing surgery at the University Medical Center Mannheim with appropriate ethical approval (approval number 2011-380N-MA). Informed consent was obtained from all participants prior to sample collection.

## Abbreviations

BET: bromodomain and extraterminal
pre-RC: pre-replication complex
HGSC: high-grade serous ovarian carcinoma
CDC7: cell division cycle 7
MCM: minichromosome maintenance
R-loop: RNA-DNA hybrid
DSB: double-strand break
HR: homologous recombination

---

**Supplementary Figure 1:**
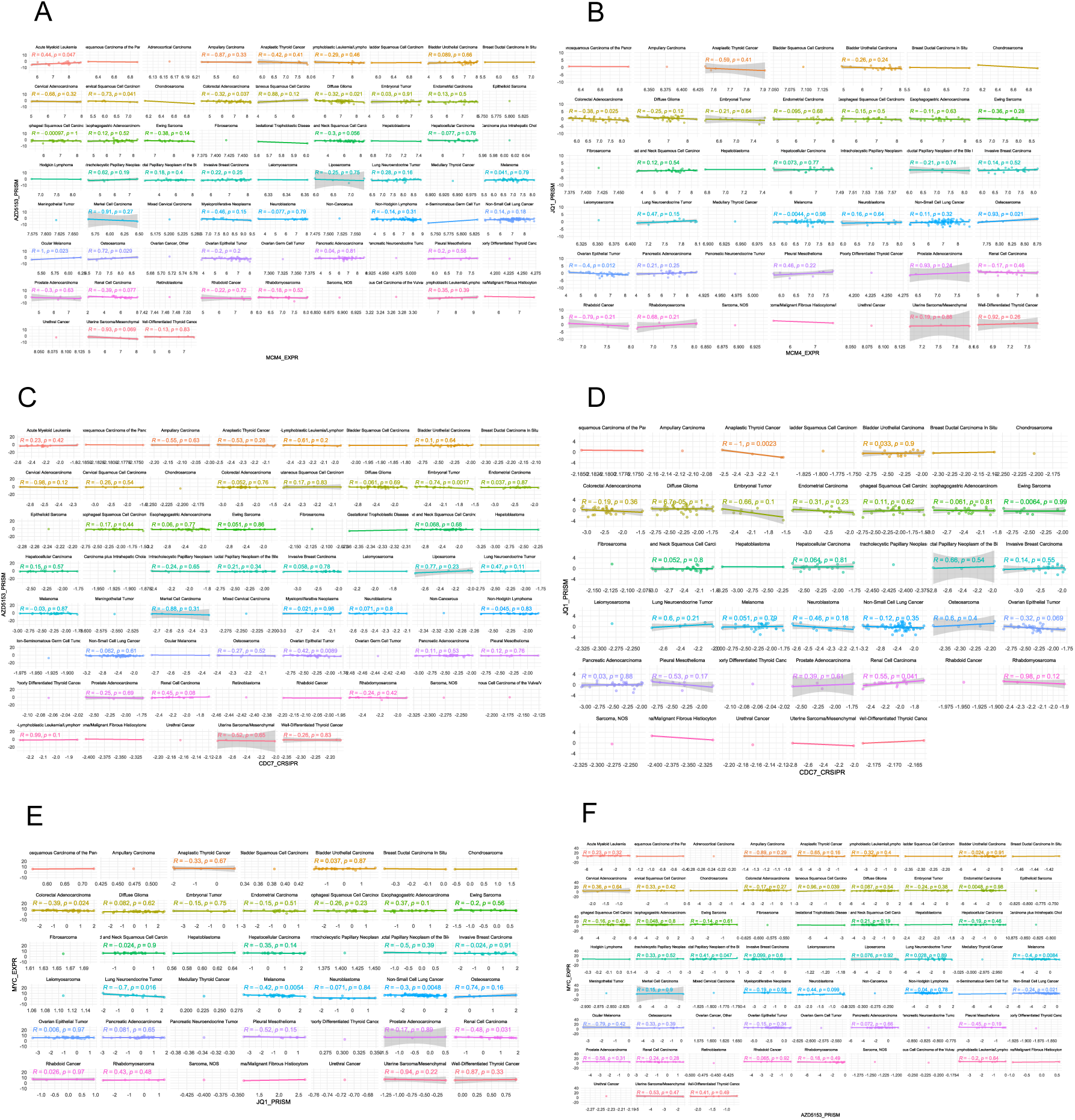
Correlation analysis of different cancer types from the CCLE and the PRISM dataset. A) Cell lines were analyzed for the dependency score of AZD5153 and MCM4 expression. B) Cell lines were analyzed for the dependency score of JQ1 and MCM4 expression.Cell lines were analyzed for the dependency score of C) Cell lines were analyzed for the dependency score of AZD5153 and CDC7_CRISPR. D) Cell lines were analyzed for the dependency score of JQ1 and CDC7_CRISPR. E) Cell lines were analyzed for the dependency score of JQ1 and MYC expression. F) Cell lines were analyzed for the dependency score of AZD5153 and MYC expression.

**Supplementary Table S1.**

| Antibody | Company | Number |
| --- | --- | --- |
| S9.6 | Sigma | MABE1095 |
| BrdU/CldU | Abcam | ab6326 |
| BrdU/IdU | Becton Dickinson | 347580 |
| CDT1 | CellSignaling | 8064S |
| MCM7 | CellSignaling | 3735S |
| GAPDH | Invitrogen | 14-9523-80 |
| pMCM7 | Sigma | PLA0034 |
| PAX8 | CellSignaling | 59019 |
| EPCAM | Invitrogen | PA5-79207 |
| P53 | CellSignaling | 48818S |
| pyH2AX | Abcam | AB22551 |
| RAD51 | Abcam | ab63801 |
| 53BP1 | CellSignaling | 88439S |
| pChk1 | CellSignaling | 2348T |
| pChk2 | CellSignaling | 2197S |
| pATR | CellSignaling | 2853S |
| MCM2 | Invitrogen | PA5-32484 |
| anti RAT Cy3 | Abcam | ab98416 |
| anti Mouse Alexa 488 | Invitrogen | A11001 |
| Anti-Rabbit Alexa 488 | Invitrogen | A11034 |
| Anti-Mouse Alexa 594 | Invitrogen | A11005 |
| Anti-Rabbit Alexa 594 | Invitrogen | A11037 |
| Goat anti Rabbit POX | Invitrogen | 31466 |
| Goat anti Mouse POX | Invitrogen | 31431 |

**Supplementary Figure 2:**
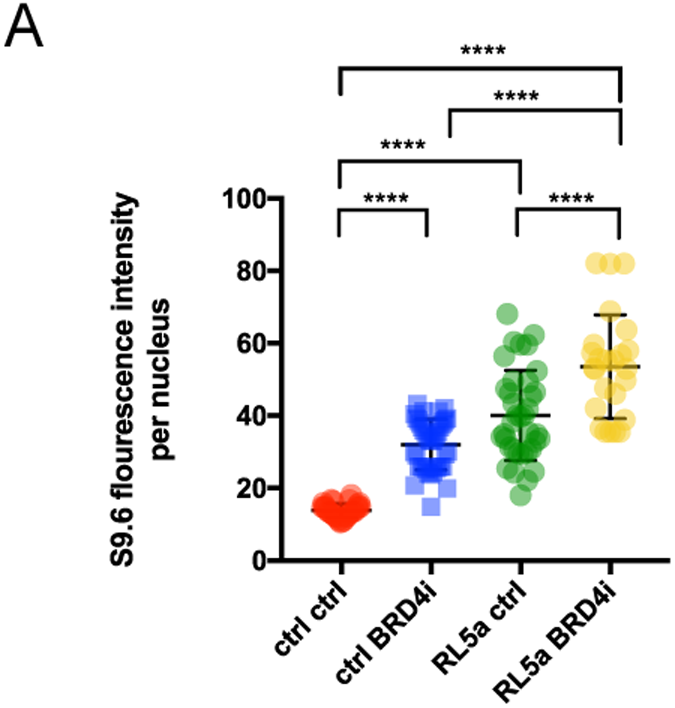
S9.6 intensity following RL5a and BRD4i treatment. A) S9.6 Immunofluorescence analysis of OVCAR8 cells treated with RL5a followed by BRD4i treatment.

**Supplementary Table S2.**
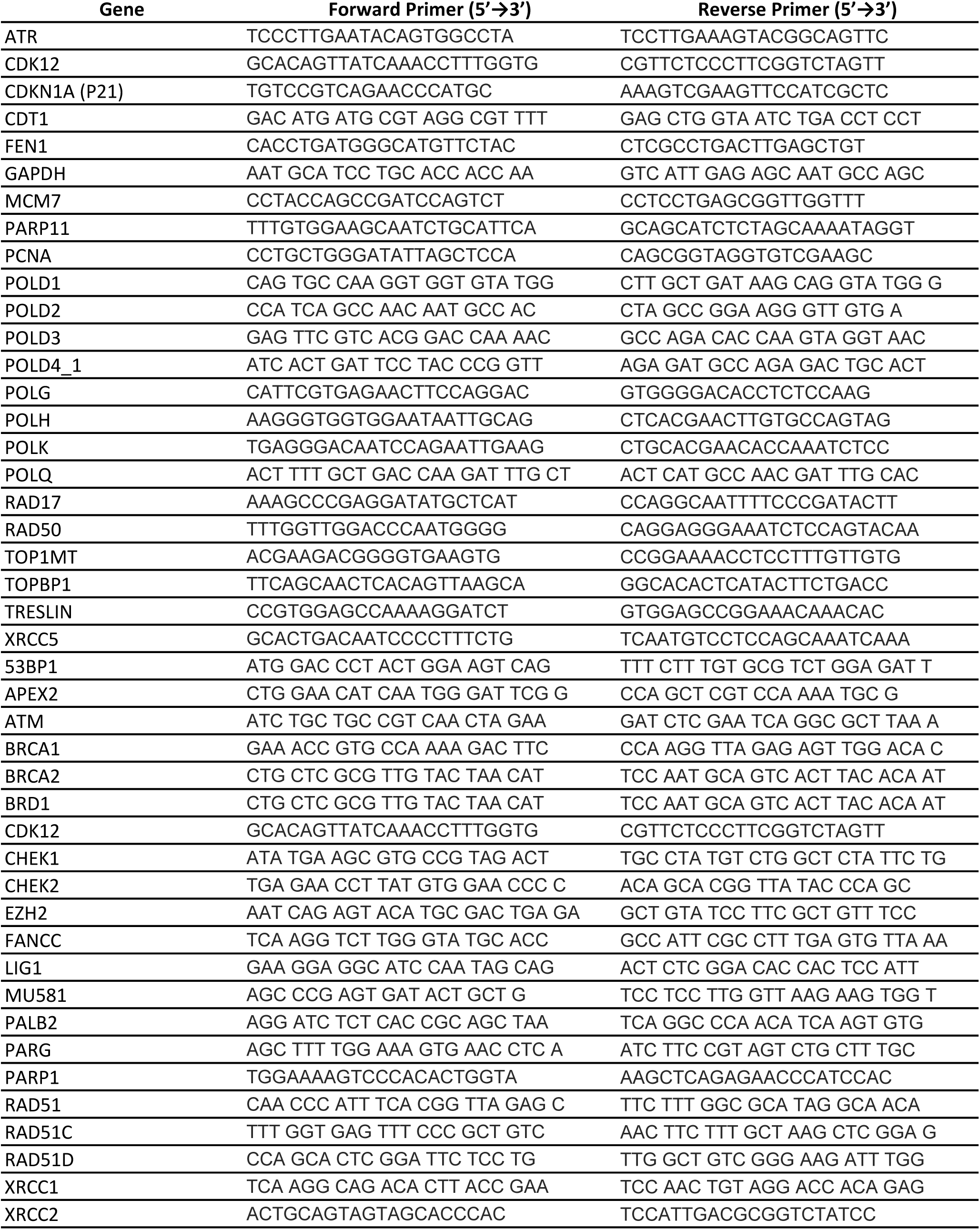

